# TRIM32 controls timely cell cycle exit in muscular differentiation through c-Myc down-regulation

**DOI:** 10.1101/2025.06.25.661523

**Authors:** Lu Xiong, Elisa Lazzari, Sabrina Pacor, Simeone Dal Monego, Erica Piovesan, Danilo Licastro, Germana Meroni

**Affiliations:** Department of Life Sciences, University of Trieste, 34127 Trieste, Italy; AREA Science Park, 34149 Padriciano, Trieste, Italy

## Abstract

Mutations in *TRIM32* cause Limb-Girdle Muscular Dystrophy recessive 8 (LGMDR8), a neuromuscular disorder primarily affecting the proximal muscles of hips and shoulders. However, the precise pathogenic mechanism remains unclear. In this study, we used *Trim32* knockout C2C12 murine myoblasts to investigate the impact of *Trim32* loss on myogenesis. We found that *Trim32* deficiency leads to impaired myogenic signaling and reduced expression of key myogenic regulatory factors ultimately leading to delayed and abnormal myotube formation. Further, we discovered that absence of Trim32 disrupts the transition from proliferation to differentiation by limiting the necessary down-regulation of the proto-oncogene c-Myc, thus delaying and altering the onset of differentation. Interestingly, unlike previous reports that emphasized protein-level regulation, our data reveal that Trim32 regulates c-Myc at mRNA stability level. Contrasting the sustained c-Myc level is able to partially recover the myogenesis defects observed in the absence of Trim32, suggesting that the Trim32–c-Myc axis may represent a promising therapeutic target for treating LGMDR8. Altogether these findings underscore the essential role of Trim32 in muscle regeneration within LGMDR8 pathogenesis.

## Introduction

Tripartite motif-containing protein 32 (*TRIM32*) was originally identified as a zinc finger protein interacting with the activation domain of lentiviral Tat proteins (Fridell *et al*, 1995). Structurally, TRIM32 contains a conserved N-terminal module, the tripartite motif, comprising a RING domain, a single type 2 B-box domain, and a Coiled-Coil region, while its C-terminal portion features a six-bladed β-propeller NHL domain (Reymond *et al*, 2001). The RING domain is responsible for the E3 ligase catalytic function (Budhidarmo *et al*, 2012; Joazeiro & Weissman, 2000) and indeed TRIM32 is primarily known for its function as an E3 ubiquitin ligase (Napolitano *et al*, 2011). Within the tripartite motif, the precise role of the B-box domain remains unclear, although some studies suggest that it modulates the rate of ubiquitin chain assembly (Lazzari *et al*, 2019), while the Coiled-Coil domain facilitates, together with the RING domain, TRIM32 homo-oligomerization which is essential for its activity (Koliopoulos *et al*, 2016). The NHL domain plays a crucial role in protein–protein interactions, likely conferring substrate specificity (Tocchini & Ciosk, 2015). Additionally, the NHL domain has also been shown to interact with RNA and growing evidence revealed that, in addition to its E3 ligase activity, TRIM32 is implicated in RNA-mediated regulation (Connacher & Goldstrohm, 2021; Kumari *et al*, 2018; Neumuller *et al*, 2008; Schwamborn *et al*, 2009).

Mutations in *TRIM32* result in an inherited rare neuromuscular disorder, Limb-Girdle Muscular Dystrophy recessive 8 (LGMDR8) (Frosk *et al*, 2002; Weiler *et al*, 1997; Weiler *et al*, 1998). This disease is characterized by the progressive wasting of muscles, mainly manifesting in the proximal muscles of the arms and legs (Caputo & Schoser, 2024). LGMDR8-associated mutations are distributed across different domains of TRIM32, with the NHL domain being the most commonly affected (Kumarasinghe *et al*, 2021). Additionally, cases with full deletion of TRIM32 have been reported (Guan *et al*, 2023; Neri *et al*, 2013). Patients carrying mutations in the NHL domain exhibit an abnormal low level of TRIM32 in muscle tissue, likely due to decreased TRIM32 protein stability (Liang *et al*, 2025; Servian-Morilla *et al*, 2019). Knock-in mice carrying an LGMDR8 corresponding pathogenic mutation (p.Asp489Asn) also presented reduced levels of mutated Trim32 protein but levels of RNA expression comparable to wild type (WT) animals, further suggesting that mutations in Trim32 alter its protein stability. Both the knock-in and full *Trim32* knock-out animal models presented a mild myopathy phenotype and both recapitulate the human phenotype (Kudryashova *et al*, 2011; Kudryashova *et al*, 2009). This evidence strongly suggests that LGMDR8 is caused by the complete loss of TRIM32 function.

LGMDR8 pathogenesis is not totally unraveled and reported findings support a role for TRIM32 in both muscle atrophy and regeneration. Early studies primarily focused on the role of TRIM32 in muscle atrophy, identifying several sarcomeric structural proteins, including actin, α-actinin, tropomyosin, desmin, and dysbindin, as potential substrates of TRIM32 E3 ubiquitin ligase activity assessed *in vitro* (Cohen *et al*, 2012; Kudryashova *et al*, 2005; Locke *et al*, 2009). However, given the low endogenous levels of TRIM32 in skeletal muscle and the highly ordered organization of these structural proteins *in vivo*, which may hinder their availability as natural substrates, it remains unclear whether TRIM32 plays a fundamental role in promoting atrophy. On the other hand, increasing evidence from both patients-derived samples and *Trim32*-deficient mouse models suggests that TRIM32 is critically involved in muscle regeneration, indicating that its role in LGMDR8 pathogenesis may extend beyond sarcomeric protein degradation and involve broader regulation of muscle homeostasis (Kudryashova *et al*, 2012; Kudryashova *et al*., 2009; Nicklas *et al*, 2012; Servian-Morilla *et al*., 2019). Myogenesis is a highly complex physiological process involving extensive crosstalk between multiple signaling pathways that collectively regulate myoblast proliferation, fusion, and differentiation (Feng *et al*, 2024; Kablar *et al*, 1998; Serrano *et al*, 2011; Smith *et al*, 1994). This process relies on the precise temporal regulation of the expression of myogenic proteins, such as MyoD, Myogenin, and Myosin Heavy Chains, and any alteration of their expression timing can result in defective muscle regeneration (Hernandez-Hernandez *et al*, 2017; Kablar *et al*., 1998; Megeney *et al*, 1996; Zammit, 2017). Notably, TRIM32 is expressed at higher levels in myoblasts than in mature skeletal muscle tissue and its expression is significantly increased during muscle regeneration after injury (Kudryashova *et al*., 2012; Kudryashova *et al*., 2011; Nicklas *et al*., 2012). These observations prompted extensive research into the role of TRIM32 in myogenesis. Indeed, several studies have highlighted a key role for TRIM32 in satellite cells biology, and TRIM32-mediated degradation of substrates such as NDRG2 and PIAS4 was shown to occur during satellite cells differentiation at the time of muscle regeneration (Kudryashova *et al*., 2012; Mokhonova *et al*, 2015).

Despite the increasing body of evidence placing TRIM32 as a central regulator of myogenesis, it is still poorly understood how its mutations may affect this process resulting in LGMDR8 (Jeong *et al*, 2023; Kumarasinghe *et al*., 2021; Shieh *et al*, 2011). To date very few studies have systematically characterized the role of TRIM32 in the myogenic process (Nicklas *et al*., 2012; Servian-Morilla *et al*., 2019). In this study, we employed *Trim32* knock-out C2C12 myoblasts to comprehensively characterize myogenesis upon loss of Trim32. We found that absence of Trim32 dramatically impairs C2C12 myogenesis, inhibiting the expression of key myogenic proteins. This impairment originates at the onset of differentiation, at which time *Trim32* knock-out cells were unable to properly exit from cell cycle, likely due to inefficient down-regulation of the pro-proliferative c-Myc transcript.

## Results

### Generation and characterization of Trim32 KO C2C12 clones

To investigate the role of Trim32 in muscular differentiation, the immortalized mouse myoblast cell line C2C12 was selected for its robust myogenic capacity (Delgado *et al*, 2003; Yaffe & Saxel, 1977). Under high-serum conditions, these myoblasts proliferate effectively, while in low-serum conditions, they start the myogenic process, differentiating and fusing into myotubes (Lawson & Purslow, 2000).

We generated *Trim32* knock-out (*Trim32* KO) C2C12 myoblasts using CRISPR/Cas9- mediated gene editing. Sequencing analysis of isolated clones revealed a recurrent c.49delG mutation in many clones as well as other inactivating *Trim32* mutations (**Fig. EV1A** and **EV1B**). Three independent single-cell KO clones were selected for subsequent experiments: two homozygous (c.49delG; p.E17Kfs*1) and one compound heterozygous (c.22_61del and c.49_50insG; p.S8Vfs*52 and p.E17Gfs*12) (**Fig. EV1B** and **Table 1**). All these mutations result in frameshifts generating premature stop codons and leading to the expression of non-functional very short N-terminal truncated Trim32 protein, if any (**Fig. 1A**). The various mutations of *Trim32* provide a reliable loss-of-function model for a deeper understanding of mechanisms underlying LGMDR8. Given that C2C12 cells differentiation potential could be affected by prolonged subculturing during gene editing procedure, we opted to use 3 *Trim32* wild-type (WT) clones, besides the original parental C2C12 cells, as control for all subsequent analyses; these clones underwent the same editing and selection protocol as the KO but resulted wild-type for *Trim32* (WT) (**Table 1**).

**Figure 1.**
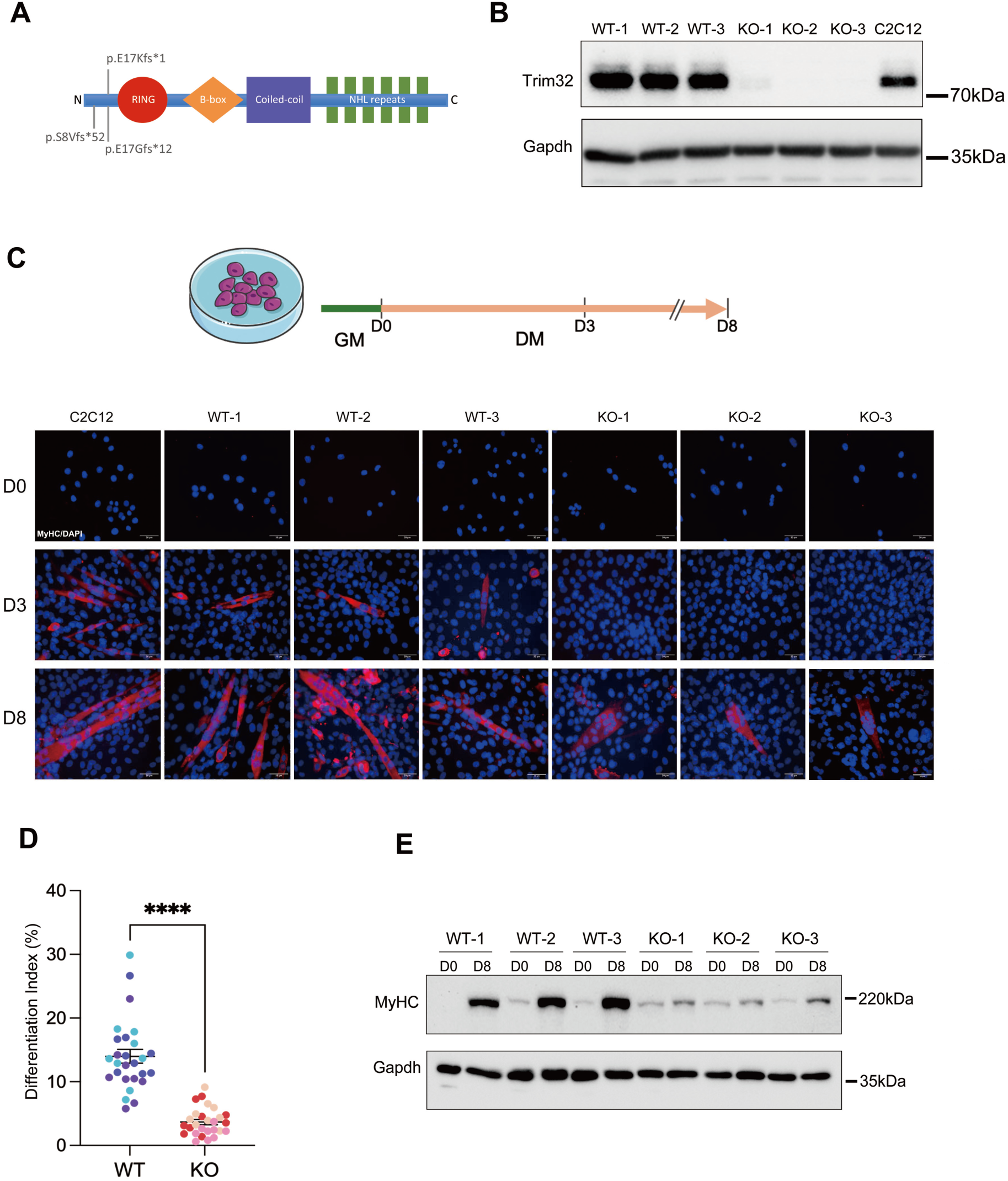
Myogenesis is impaired in *Trim32* KO cells. **(A)** Scheme of the *Trim32* mutations generated in the C2C12 clones and resulting in premature stop codons and leading to non-functional fragments: p.E17Kfs*1, p.S8Vfs*52 and p.E17Gfs*12. **(B)** Representative Western blot of Trim32 in parental C2C12 cells, three KO clones, and three WT clones (n=3). Trim32 protein was detected using an antibody against its N-terminus. **(C)** Top, scheme of the differentiation timeline adopted. Bottom, immunostaining for MyHC (red) at day 0 (D0), and at 3 (D3) and 8 days (D8) of differentiation, in parental C2C12 cells, 3 WT clones and 3 *Trim32* KO clones. Nuclei were counterstained with DAPI (blue) (Scale bar = 50 µm; magnification 40x; n = 3). (**D)** Differentiation index of *Trim32* KO and WT clones at D8. The differentiation index is calculated as the ratio of nuclei within MyHC-positive myotubes to the total number of nuclei (mean ± SD; unpaired t-test, ****P < 0.0001; n = 3). Each dot represents a microscope field; shades of blue and red represent the different clones **(E)** Representative Western blot of MyHC protein levels in *Trim32* KO and WT clones at D0 and D8 of differentiation as indicated (n = 3). Gapdh was used as loading control.

**Table 1.** Genotypes of the generated C2C12 clones.

|  | Clones | *Trim32* Genotype | *Trim32* Mutations |
| --- | --- | --- | --- |
| KO | KO-1 | c.49delG/c.49delG | p.E17Kfs*1 |
|  | KO-2 | c.49delG/c.49delG | p.E17Kfs*1 |
|  | KO-3 | c.22_61del/c.49_50insG | p.S8Vfs*52/p.E17Gfs*12 |
| WT | WT-1 | WT | none |
|  | WT-2 | WT | none |
|  | WT-3 | WT | none |

The absence of the Trim32 protein in KO clones was confirmed with Western blot analysis using two polyclonal antibodies targeting either the N-terminal (**Fig. 1B**) or the central regions (**Fig. EV1C**) of Trim32.

### Absence of Trim32 impairs terminal differentiation of C2C12 cells

Having generated *Trim32* KO C2C12 clones, we started assessing the differentiation of these cells in culture. In the classic differentiation protocol, C2C12 cells are grown in proliferating medium (Growth Medium, GM) and then triggered to differentiate upon serum reduction (Differentiation Medium, DM) (see Methods). A myosin heavy chain subtype, Myosin Heavy Chain 3 (MyHC), was used as a marker to monitor differentiation. MyHC is a structural protein of the sarcomere and is exclusively expressed in differentiating myocytes and myotubes (Schiaffino, 2018; Schiaffino *et al*, 2015). Parental C2C12 cells, along with three WT and three KO clones, were seeded in GM and allowed to proliferate for 24 hours before switching to DM to induce differentiation up to 8 days.

Parental C2C12 and WT cells initiated the myogenic program promptly under differentiation conditions. By day 3 (D3), immunofluorescence images showed MyHC-positive staining in differentiating fusiform myocytes (mononuclear) and small myotubes (containing two or very few nuclei). In contrast, KO clones exhibited remarkably reduced myogenic potential. No MyHC-positive cells were detected in *Trim32* KO clones after 3 days of differentiation, indicating that they were unable to undergo proper myogenic progression (**Fig. 1C**). By day 8 (D8), WT clones showed formation of myotubes to a similar extent as parental C2C12, while KO clones appeared severely impaired in their ability to generate myotubes. Additionally, the myotubes formed by *Trim32* KO clones were significantly smaller, exhibited clustered nuclei, and lacked the proper elongated, stretched morphology observed in WT myotubes (**Fig 1C**).

The aberrant differentiation of KO clones was confirmed by determining the differentiation index: compared to an average of 13.99% of nuclei within MyHC-positive myotubes in WT clones, the percentage was only 3.68% in KO cells after full-term differentiation (D8) (**Fig. 1D**). Western blot analysis of MyHC expression during differentiation further supported these findings, revealing a substantially lower MyHC level in KO clones after 8 days of differentiation (**Fig. 1E**). This reduction was consistent with the observed defect in myotube formation in KO cells.

These results indicate the critical requirement of Trim32 for C2C12 myoblast myogenesis and suggest that deletion of *Trim32* causes myogenic defects without completely eliminating myotube formation in C2C12 cells.

### Transcriptomic analysis of Trim32 WT and KO clones under proliferation and differentiation conditions

From the previous observations, it is evident that the absence of Trim32 has a great impact already at 3 days of C2C12 differentiation. In parallel, we conducted an unbiased transcriptomic profiling to identify early global gene expression changes influenced by the *Trim32* genotype during proliferation (D0) and initial differentiation (D3) stages (**Fig. 2A**). Multidimensional Scaling (MDS) confirmed the segregation of gene expression profiles based on *Trim32* genotype (Dim2, 16% variance: WT vs. KO) and differentiation timepoint (Dim1, 44% variance: D0 vs. D3) (**Fig. 2A**). Similarly, the cluster dendrogram classified WT and KO clones into distinct groups at both D0 and D3, reflecting their different gene expression patterns (**Fig. 2B**). On day 3, *Trim32* WT and KO cells were transcriptionally more distinct compared to day 0 as indicated by the increased distance along the Dim 2 axis at D3 (**Fig. 2A**). This suggests that the influence of *Trim32* becomes progressively more significant during the process of myogenic differentiation.

**Figure 2.**
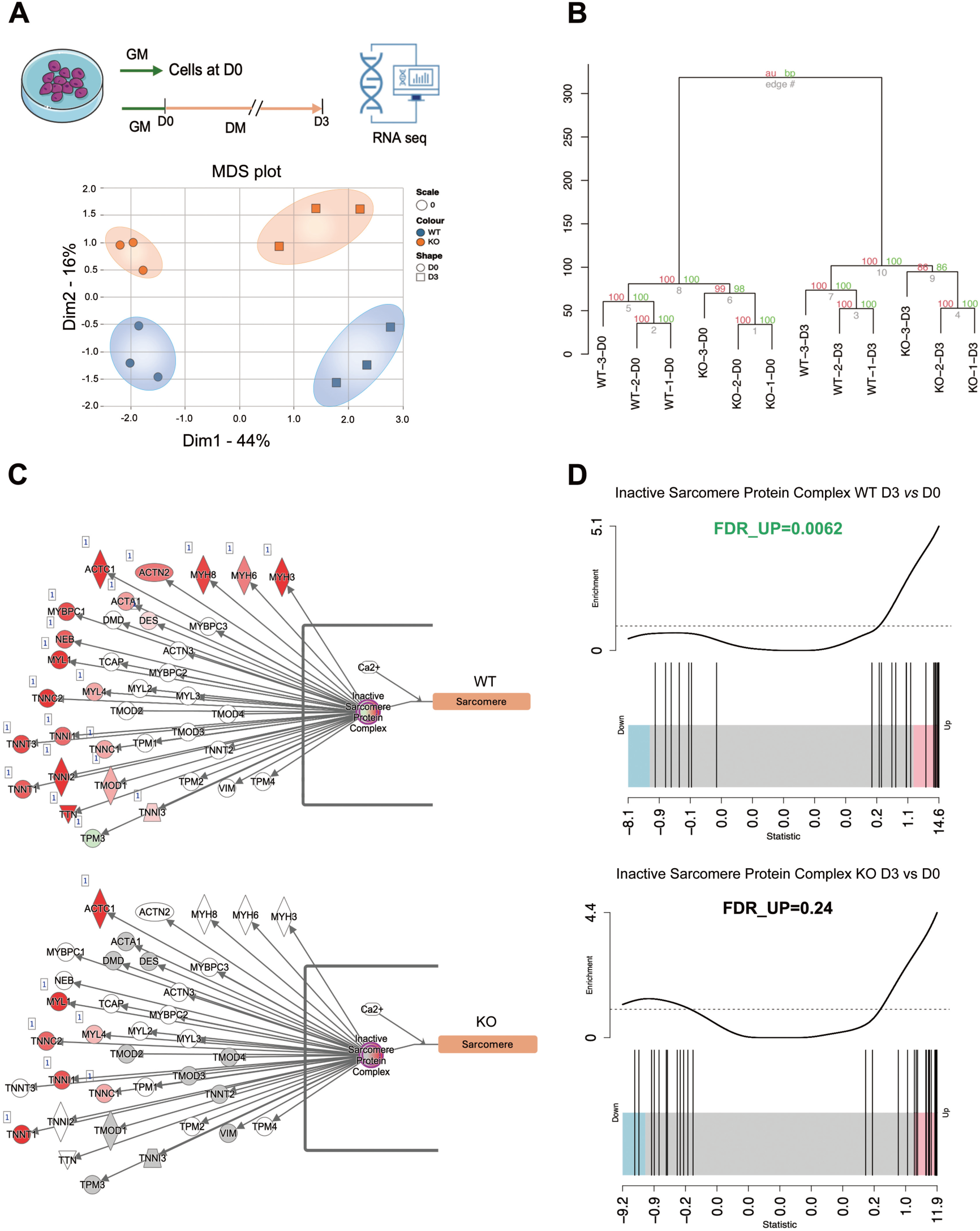
Transcriptomic analysis reveals defective myogenesis program in *Trim32* KO cells at Day 3. **(A)** Scheme of differentiation for the RNAseq experiment and MDS analysis of RNAseq data from three *Trim32* KO and three WT clones in proliferation (D0) and differentiation (D3). Dim1: D0 *vs* D3; Dim2: *Trim32* WT *vs* KO clones. Genotypes and time points are indicated in the graph with color and shape code, respectively. (**B)** Cluster dendrogram grouping WT and *Trim32* KO clones at day 0 (D0) and day 3 (D3) of differentiation. **(C)** Genes belonging to the Inactive Sarcomere Protein Complex pathway activated from D0 to D3 (red symbols) in WT (upper scheme) and *Trim32* KO (lower scheme). **(D)** Gene set enrichment barcode plots show the expression change of each gene within the Inactive Sarcomere Protein Complex pathway after 3 days of differentiation in WT (FDR_UP=0.0062) and *Trim32* KO (FDR_UP=0.24).

Pathway analysis of the differentially expressed transcripts identified, as significantly enriched, the class of genes associated with the “inactive sarcomere protein complex” pathway. Key transcripts are significantly upregulated in WT clones at D3 compared to D0, including different subtypes of Myosin Heavy Chains (*MYH3*, *MYH6*, *MYH8*), α-Actin 1 (*ACTA1*), α-Actinin 2 (*ACTN2*), Desmin (*DES*), Tropomodulin 1 (*TMOD1*), and Titin (*TTN*), consistently with what reported in literature. On the contrary, most of the up-regulated transcripts in WT are either not detected in the KO clones after 3 days of differentiation or they are up-regulated to a lower extent (**Fig. 2C**). Gene set enrichment bar-code plots for the same network also confirmed the enrichment of genes in WT clones at the early stages of differentiation (FDR_UP = 0.0062) whereas this is not the case in *Trim32* KO clones (FDR_UP = 0.24) (**Fig. 2D**). Since many of these genes encode major structural proteins of the sarcomere, the smallest contractile unit within the cytoplasm of skeletal muscle fibers, these data indicate that absence of Trim32 results in molecular impairment of muscular differentiation.

These findings further support our initial data that the absence of Trim32 impairs the ability of C2C12 cells to undergo proper myogenesis already at early stages of the process.

### Trim32 deficiency dysregulates MyoD and Myogenin expression

Myogenesis is a tightly regulated process that is achieved by controlling the expression of Myogenic Regulatory Factors (MRFs) specifically in muscle cells and within precise time windows corresponding to different stages (Asfour *et al*, 2018). Among these MRFs, MyoD and Myogenin play critical roles ultimately regulating MyHC expression. Given the impaired myogenesis and significantly perturbed early differentiation markers in *Trim32* KO clones, we decided to analyze these two upstream regulators of MyHC.

MyoD, also known as myoblast determination protein 1, forms heterodimers with basic helix-loop-helix (bHLH) E-proteins to recognize and regulate the transcription of myogenic target genes. MyoD activation occurs in myoblasts prior to the onset of differentiation, underscoring its role as an early regulator of myogenesis (Tapscott, 2005). Normalized expression from the RNAseq analysis revealed, despite high variability, lower transcriptional levels of *MyoD* in *Trim32* KO clones compared to WT clones during both proliferation (D0) and differentiation (D3) (**Fig. 3A**). Given MyoD high expression during myoblasts proliferation, we checked its protein levels at D0. Consistent with a reduced myogenic potential, *Trim32* KO clones exhibited significantly lower MyoD protein levels compared to the WT clones (**Fig. 3B**).

**Figure 3.**
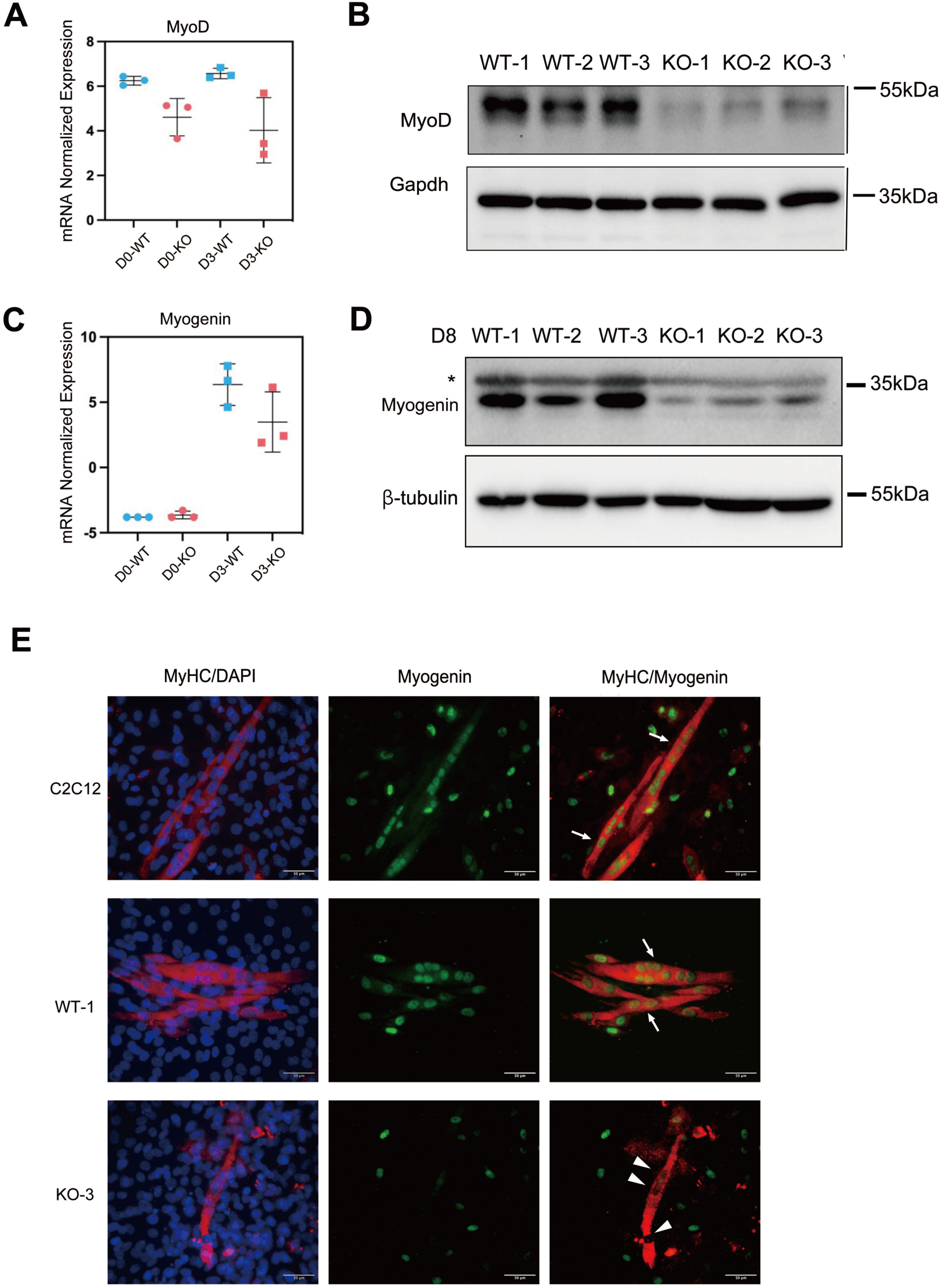
The absence of *Trim32* affects Myogenic Regulatory Factors expression. **(A)** Normalized *MyoD* transcript levels in WT and *Trim32* KO clones at 3 days of differentiation (mean ± SD; n = 3). **(B)** Representative Western blot analysis of MyoD protein levels in WT and *Trim32* KO clones cultured in growth medium (D0) (n = 3). (**C)** Normalized *Myogenin* transcript levels in WT and *Trim32* KO clones at 3 days of differentiation (mean ± SD; n = 3). **(D)** Representative Western blot analysis of Myogenin protein levels in WT and *Trim32* KO clones at 8 days (D8) of differentiation (n = 3). Asterisk indicates a non-specific band. **(E)** Immunostaining for MyHC (red) and Myogenin (green) of representative WT and *Trim32* KO clones, and parental C2C12 cells at 8 days of differentiation. Arrowheads indicate nuclei lacking Myogenin expression in *Trim32* KO myotubes; Arrows indicate Myogenin-positive nuclei in myotubes of WT and parental C2C12. Nuclei are counterstained with DAPI (blue) (Scale bar = 50 µm; magnification 40x; n = 3).

Myogenin is instead pivotal during later stages of myogenic differentiation. Elevated Myogenin levels are observed in myoblasts that exit the cell cycle and differentiate into myotubes (Wright *et al*, 1989). While both WT and KO clones exhibited increased *Myogenin* transcriptional expression after 3 days of differentiation, *Trim32* KO clones showed a limited increase compared to WT clones (**Fig. 3C**). In our experimental setting, Myogenin protein was detected at day 8 of differentiation, consistent with it affecting the terminal stage of myogenesis. Coherently, *Trim32* KO clones showed significant lower Myogenin protein levels compared to WT clones (**Fig. 3D**), in keeping with reduced upregulation of Myogenin transcript observed during early phases of differentiation. Immunofluorescence images further corroborated this finding, showing markedly reduced Myogenin positive cells in *Trim32* KO clones after 8 days of differentiation, consistent with reduced differentiation potential into myotubes (**Fig. 3E** and **Fig. EV2**). Interestingly, the residual Myogenin positive cells did not correspond to the few and abnormal myotubes formed by *Trim32* KO clones, whereas Myogenin was abundantly detected in the nuclei of WT and parental C2C12 myotubes (**Fig. 3E** and **Fig. EV2**). Based on this finding, we suggest that the lack of Myogenin in the initially formed KO myotubes hinders their further differentiation.

Overall, our data suggest that Trim32 deficiency diminishes MyoD and Myogenin levels, and possibly their activity from the early stages of differentiation. Perturbation of the key myogenic regulatory factors regulatory network may underscore the observed impaired myotube formation.

### Trim32 regulates cell cycle exit during the transition from proliferation to differentiation

At the onset of myogenic differentiation, a complex MRFs regulatory network induces cells to reduce their proliferation rate to eventually exit the cell cycle, a prerequisite to start the differentiation program (Dumont *et al*, 2015; Shen *et al*, 2003). We noticed that KO cells continued to proliferate even after differentiation was induced and also, cell cycle related pathways were enriched in the transcriptomic analysis (data not shown). Therefore, we further investigated the role of Trim32 in regulating cell cycle dynamics during this critical transition.

Preliminary analysis using the MTT assay indicated higher metabolic activity in *Trim32* KO clones over the course of differentiation, potentially suggesting a higher proliferation rate in KO clones compared to WT (**Fig. EV3A**). Notably, since differences in proliferation were detectable by MTT already after 2 days of differentiation, we also assessed the positivity for Phospho-histone H3 (Ser10, pHH3), as a marker of mitosis, under growth conditions (D0) and 2 days after differentiation onset (D2). At D0, similar percentages of pHH3-positive myoblasts were observed in *Trim32* KO and WT C2C12 cells, indicating comparable mitotic rates under proliferating conditions (**Fig. 4A**, upper panels and graph). Upon differentiation induction, proliferation slowed down in both *Trim32* KO and WT clones, as evidenced by a decrease in the percentage of pHH3-positive myoblasts from D0 to D2. However, at day 2, a significantly higher percentage of pHH3-positive myoblasts was observed in *Trim32* KO clones compared to WT, indicating the presence of more KO cells still undergoing mitosis (on average from 4.1% to 0.2% in WT; and from 4.3% to 1.2% in KO) (**Fig. 4A**, lower panels and graph).

**Figure 4.**
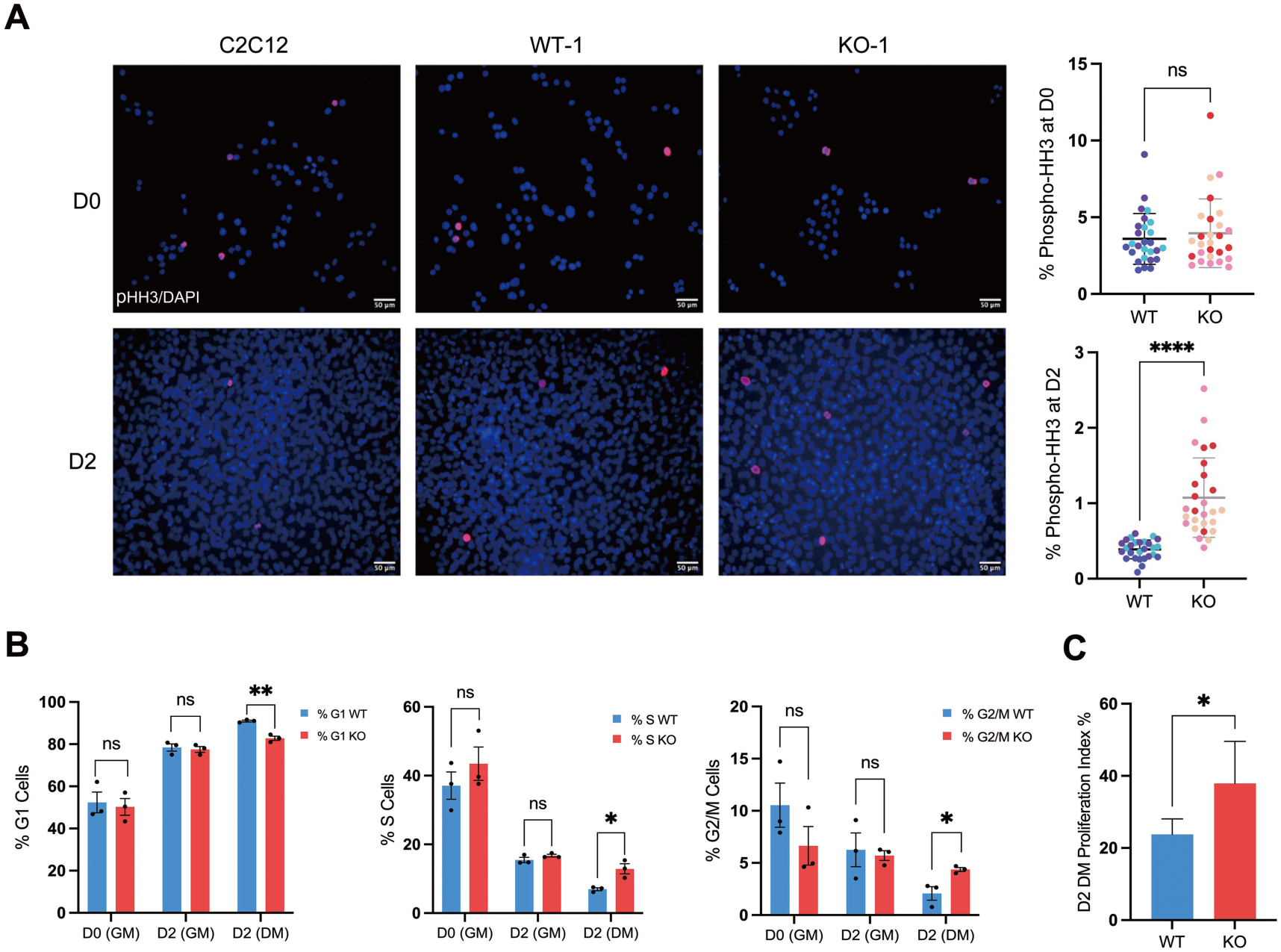
*Trim32* KO cells exhibit prolonged proliferation after the onset of differentiation. **(A)** Phospho-histone H3 (pHH3, red) immunostaining. Images of representative WT and *Trim32* KO clones, and C2C12 parental cells in proliferation (D0) and after 2 days of differentiation (D2) (Scale bar = 50 µm; magnification 20x). On the right side, graphs of the corresponding quantification; each dot represents a microscope field; shades of blue and red represent the different clones (n = 3; mean ± SD; unpaired t-test, ****P <0.0001). **(B**) Flow cytometry quantification of the percentage of cells in G1, S and G2/M phase for WT and *Trim32* KO clones in growth medium at Day 0 (D0 (GM)) and Day 2 (D2 (GM)), as well as Day 2 after the induction of myogenic differentiation (D2 (DM)) (mean ± SD; unpaired t-test, \**P<*0.1, \*\**P<*0.01, n = 3). **(C)** Graph showing the proliferation index in *Trim32* KO and WT clones using CFSE (Carboxyfluorescein Diacetate Succinimidyl Ester) 2 days after initiating differentiation (D2 DM) (mean ± SEM; n = 2; unpaired t-test, P < 0.05).

At this point, we analyzed cell cycle progression by flow cytometry and observed that, upon culturing cells in differentiation medium for 2 days, as expected the proportion of cells in the G1 phase increased, accompanied by a decrease in the proportion of cells in the S and G2/M phases. This change, observed in both WT and *Trim32* KO clones, indicates progressive cell cycle withdrawal at the start of differentiation (**Fig. 4B**). However, at D2, a lower percentage of *Trim32* KO cells was found in the G1 phase compared to WT cells, while a correspondent higher proportion of KO cells were in the S and G2/M phases (**Fig. 4B**). These differences between WT and KO clones were not observed when cells were maintained in growth medium at either D0 or for 2 days (D2, GM) (**Fig. 4B**). These findings suggest that *Trim32* KO cells are unable to efficiently withdraw from the cell cycle in response to the differentiation stimulus. The greater proportion of KO cells in the G2/M phase indicates a higher proliferation rate. To further support this, we used Carboxyfluorescein Diacetate Succinimidyl Ester (CFSE) to monitor cell division. CFSE covalently labels cytoplasmic proteins with a fluorescent dye, and during cell division, the marker is equally distributed between daughter cells, resulting in reduced fluorescence intensity. Thus, cell proliferation can be quantified by measuring the fluorescence dilution using flow cytometry. At 2 days post-induction of differentiation, *Trim32* KO clones exhibited a significantly higher proliferation index compared to WT clones further confirming the above results (**Fig. 4C** and **EV3B**).

Overall, these results indicate that Trim32 deficiency interferes with proper and timely cell cycle exit likely delaying the initiation and/or the extent of differentiation in C2C12 cells.

### Trim32 regulates c-Myc mRNA stability during cell cycle withdrawal

To further investigate the molecular mechanism underlying Trim32 role in cell cycle withdrawal during the transition from proliferation to differentiation, we examined its effect on c-Myc regulation. Previous studies suggested that Trim32 interacts with and ubiquitinates c-Myc, leading to its proteasome-mediated degradation (Hillje *et al*, 2011; Nicklas *et al*, 2019; Nicklas *et al*., 2012; Schwamborn *et al*., 2009). c-Myc proto-oncogene plays a crucial role in regulating cell fate across various cell types, including C2C12 myoblasts (Luo *et al*, 2019; Seyer *et al*, 2006). Notably, sustained c-Myc expression in C2C12 myoblasts inhibits differentiation and myotube formation (Denis *et al*, 1987; Seyer *et al*., 2006). Indeed, we observed a progressive reduction in c-Myc levels in C2C12 parental cells during the course of differentiation (**Fig. EV4A**), and the inhibition of MyHC level upon transient overexpression of c-Myc at the onset of differentiation in C2C12 (**Fig. EV4B**). Our data are in agreement with the fact that c-Myc turnover is essential for initiating differentiation (Luo et al., 2019). In both *Trim32* WT and KO clones, Western blot analysis revealed indeed significant reduction in c-Myc levels after 6 hours in differentiation medium. However, c-Myc levels remained relatively higher in KO clones compared to WT clones at this stage (**Fig. 5A**). To further consolidate the functional meaning of the observed differential expression of c-Myc, we analyzed the transcriptional levels of known c-Myc downstream target genes after 3 days of differentiation (Cairo *et al*, 2005). Among the genes activated by c-Myc, the three we tested - *Ccnb1, Rcl,* and *Tnbs1* - all showed higher levels in KO clones; accordingly, three of the genes inhibited by c-Myc - *Adm*, *Eca40,* and *Tert* - exhibited lower transcript levels in KO clones, indicating that the observed higher levels of c-Myc correlate with its elevated activity (**Fig. 5B**).

**Figure 5.**
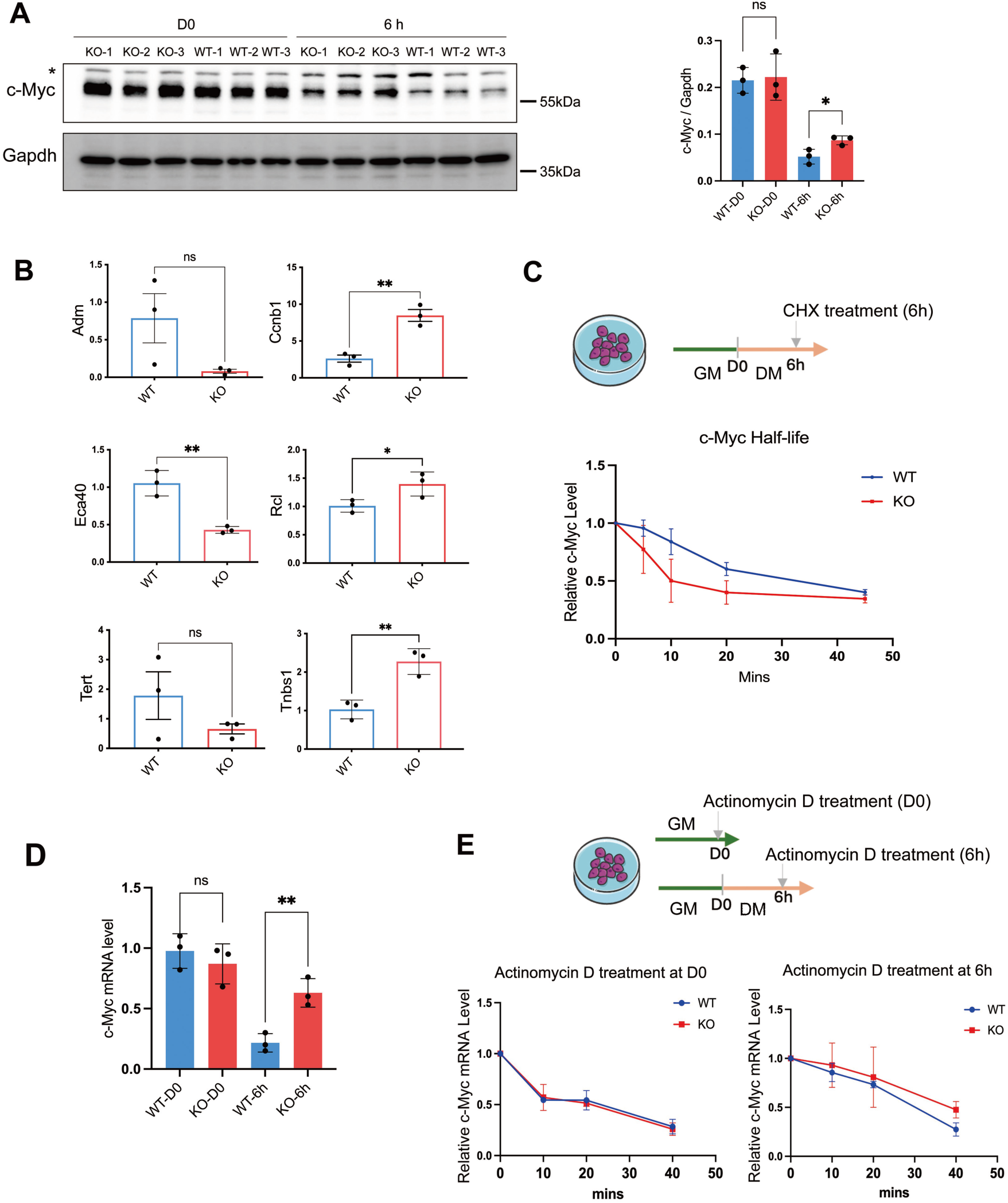
Trim32 regulates c-Myc level via mRNA stability. **(A)** Representative Western Blot analysis of c-Myc in *Trim32* KO and WT clones at D0 (in GM) and 6h after the induction of myogenic differentiation. Asterisk indicates a non-specific band. On the right side, graph of the corresponding quantification (mean ± SD; n=3; unpaired t-test, *P<0.1). **(B)** Transcriptional changes of c-Myc downstream target genes in WT and *Trim32* KO clones analyzed by real-time RT-PCR after 3 days of differentiation (mean ± SD; n = 3, unpaired t-test, *P<0.1, **P<0.01). **(C)** Graph showing the stability of c-Myc protein in WT and *Trim32* KO clones: cells were cultured in DM for 6h hours before treatment with Cycloheximide (CHX) for the indicated time points and c-Myc level was detected by immunobloting (mean ± SD; n = 3). **(D)** Real-time RT-PCR analysis of c-Myc mRNA steady state level in WT and *Trim32* KO clones at D0 and at 6 hours of differentiation (6h) (mean ± SEM; n = 3). **(E)** Stability of c-Myc mRNA determined by Actinomycin D chase in WT and *Trim32* KO cells, cultured either in GM or DM for 6h, for the indicated time points. Relative RNA levels were measured by real-time RT-PCR (c-Myc primers-1) normalized to Gapdh level (mean ± SD; n = 3).

Based on this evidence and given that c-Myc has been reported as a potential substrate of Trim32, we hypothesized that Trim32 could promote C2C12 cell cycle withdrawal by facilitating c-Myc degradation. We therefore examined its stability at the onset of differentiation using pulse-chase with the Cycloheximide (CHX) protein synthesis inhibitor. Unexpectedly, c-Myc half-life resulted not extended in *Trim32* KO clones when compared to WT, suggesting that Trim32 does not facilitate c-Myc degradation in this context (**Fig. 5C**). Furthermore, subcellular fractionation revealed that c-Myc and Trim32 localize to different cellular compartments, i.e. nucleus and cytoplasm, respectively (**Fig. EV4C**). Consistently, immunoprecipitation experiments performed after 6 hours of differentiation failed to detect a direct interaction between Trim32 and c-Myc (**Fig. EV4D**). Together, these findings suggest that the increased c-Myc levels detected in *Trim32* KO clones at the onset of differentiation are unlikely due to enhanced protein stability resulting from Trim32 ablation.

We therefore hypothesized that Trim32 might regulate c-myc abundance by acting at transcriptional level. Interestingly, real-time RT-PCR analysis showed that while c-Myc mRNA levels were not affected by Trim32 in growth conditions, the absence of Trim32 was associated with a higher c-Myc mRNA level compared to WT cells observed 6 hours after switching to DM, i.e. the physiological WT reduction was not observed in KO cells (**Fig. 5D**). Given the growing role of TRIM proteins in regulating mRNA stability (Connacher & Goldstrohm, 2021), we used Actinomycin D pulse treatment to inhibit transcription and measure c-Myc mRNA half-life. Trim32 has no effect on the stability of c-Myc mRNA in proliferating conditions, but upon induction of differentiation the stability of c-Myc mRNA resulted enhanced in *Trim32* KO clones (**Fig. 5E** and **Fig. EV4E** and **EV4F**). The stability of the c-Fos mRNA, analyzed as control, was not affected by Trim32 in either condition (**Fig. EV4G**). The enhanced stability of c-Myc transcript at the onset of differentiation can explain the elevated levels of c-Myc protein observed in KO clones despite no change, or even reduction, of protein stability in these samples, and may correlate with delayed cell cycle exit observed in the absence of Trim32.

In summary, these findings suggest that Trim32 regulates c-Myc levels primarily by mRNA stabilization that results in persistent protein level at the onset of the differentiation stimulus, rather than through direct interaction or protein degradation. We propose that this regulation might be essential for proper cell cycle exit and subsequent myogenic differentiation.

### Counteracting c-Myc sustained expression partially rescues Trim32 KO defects

Sustained c-Myc levels observed in *Trim32* KO clones may influence the transition from proliferation to differentiation. To further validate the involvement of the Trim32-c-Myc axis in differentiation, we analyzed MyoD and Myogenin expression in *Trim32* KO upon c-Myc silencing in an attempt to rescue the differentiation defect observed in the KO clones. c-Myc down-regulation was achieved by transfecting siRNA into proliferating KO cells, and silencing was validated via real-time RT-PCR and Western blot after 24 hours in growth medium (D0) (**Fig. 6A**, upper panel and **6B**, left graph). No significant difference in MyoD transcript was observed after 6 hours of differentiation (data not shown), suggesting that MyoD expression is not regulated by c-Myc at this stage and consistently with the observed decrease of MyoD before the switch (**Fig. 3B**). Of note, downregulation of c-Myc in the KO clones was sufficient to elevate Myogenin RNA to levels detected in WT clones transfected with si-Control, i.e. normal levels, at D3 of differentiation (**Fig. 6B**, right graph). This transcriptional increase resulted in a full rescue at the protein level in KO cells at later stages of differentiation (D8) (**Fig. 6A**, lower panel). Immunostaining further revealed that downregulation of c-Myc in *Trim32* KO clones can increase Myogenin levels and restore its expression also in nuclei of myotubes, as for the WT cells (**Fig. 6C**). However, c-Myc down-regulation is not able to rescue KO myotube morphology to WT level, neither to significantly increase the differentiation index (data not shown).

**Figure 6.**
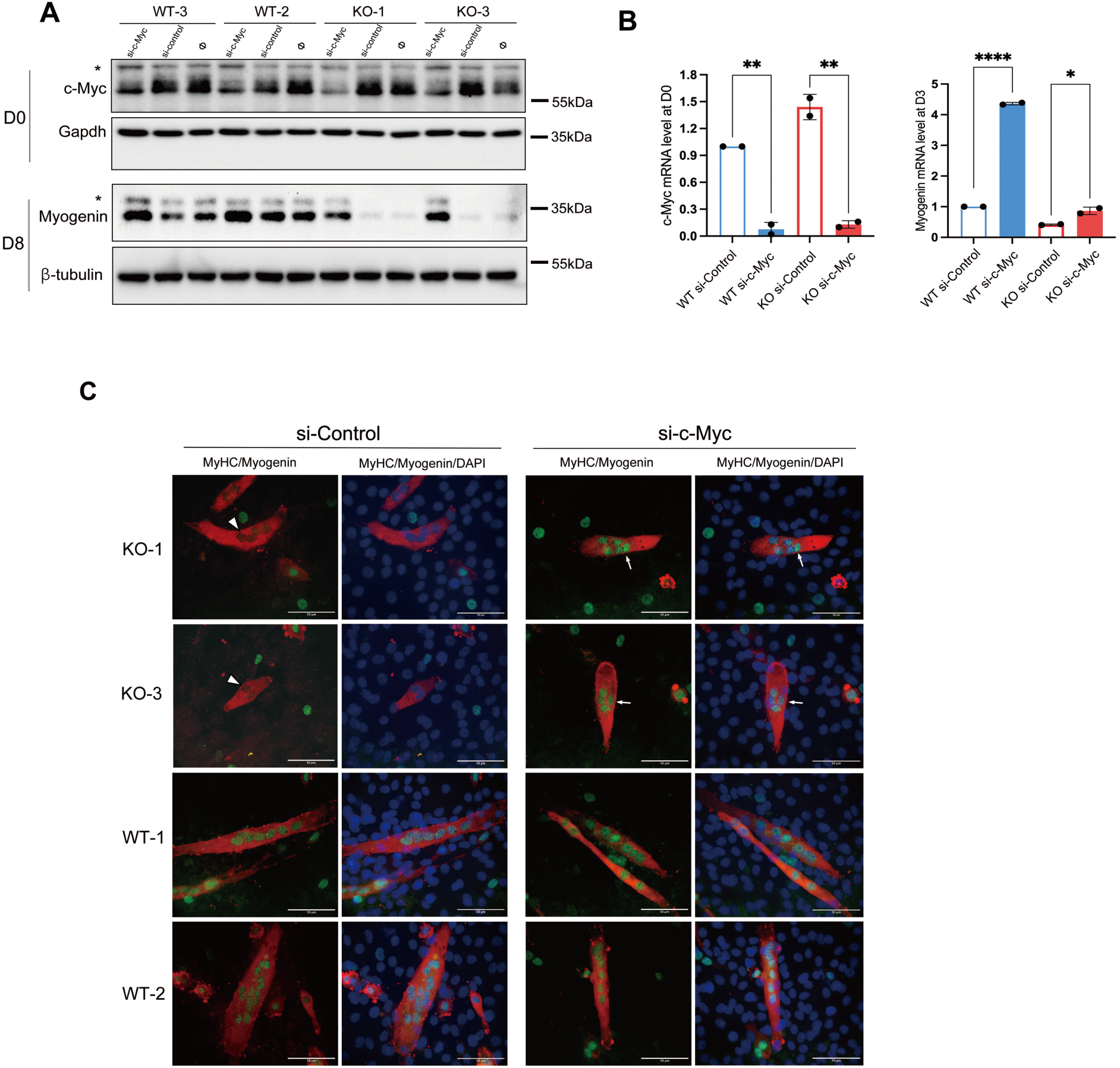
c-Myc silencing partially rescues *Trim32* KO defects. **(A)** Representative Western Blot analysis of WT and *Trim32* KO clones transfected with si-c-Myc, si-Control, and without siRNA. c-Myc levels were detected 24 hours post-transfection in GM (D0) (upper panel); Myogenin levels were detected at 8 days of differentiation (lower panel). Asterisk indicates a non-specific band. **(B)** c-Myc (left graph) and Myogenin (right graph) transcriptional levels were detected in WT and *Trim32* KO cells upon transfection with control siRNA (WT-si-con and KO-si-con) and with c-Myc siRNA (WT-si-c-Myc and KO-si-c-Myc). c-Myc mRNA was detected by real-time RT-PCR after 24 hours of culture in GM (D0), and Myogenin was detected at 3 days (D3) of differentiation (mean ± SD; unpaired t-test, *P < 0.1, **P < 0.01, ****P < 0.001; n=3). **(C)** Immunostaining for MyHC (red) and Myogenin (green) of WT and KO clones as indicated, upon silencing with control siRNA (si-Control), or c-Myc siRNA (si-c-Myc) at 8 days of differentiation. Arrowheads indicate myotubes displaying nuclei lacking Myogenin in KO-si-control, while arrows indicate Myogenin-positive nuclei in KO-si-c-Myc myotubes. Nuclei are counterstained with DAPI (blue) (Scale bar = 50 µm; magnification 63x).

Together, these findings suggest that precise c-Myc turnover is essential for effective C2C12 differentiation and that the Trim32–c-Myc regulatory axis plays a critical role in the expression of key myogenic factors, particularly Myogenin, within a narrow time window upon the differentiation stimulus.

## Discussion

Although mutations in *TRIM32* are widely recognized as the cause of LGMDR8, the precise pathogenic mechanism underlying the disease remains unclear. In this study, we used *Trim32* KO C2C12 myoblast cells to investigate the role of Trim32 in muscular homeostasis and found that its absence significantly hinders myogenesis supporting a critical involvement of Trim32 in muscle regeneration.

Our results show that the absence of Trim32 in C2C12 cells leads to impaired differentiation, characterized by the formation of fewer and smaller myotubes and lower MyHC expression. At early stages of differentiation, we further observed lower levels of expression of genes encoding sarcomeric proteins and key MRFs, such as MyoD and Myogenin. These observations are consistent with previous findings in *Trim32* full KO and D489N LGMDR8-associated mutation knock-in mice, which display smaller muscle fibers, small myofibers with centrally located nuclei and a lack of developmental MyHC-positive myotubes, indicating defective muscular regeneration (Kudryashova *et al*., 2011; Kudryashova *et al*., 2009). Formation of fewer and abnormally shaped myotubes and reduced expression of differentiation markers such as Myogenin and MyHC, was also observed in primary myoblasts from LGMDR8 patients, further confirming the crucial role of TRIM32 for proper myogenesis (Servian-Morilla *et al*., 2019).

Interestingly, in the context of C2C12 myogenesis process we also observed that Trim32 plays a role in cell proliferation. *Trim32* KO clones failed to properly exit the cell cycle upon the initiation of differentiation and maintained a higher proliferation rate compared to WT clones even 2 days post-differentiation onset. This dysregulation in the transition from proliferation to differentiation likely contributes to the observed reduced expression of Myogenin and MyHC. While Trim32 role in regulating cell proliferation has been studied extensively in cancer, its pro- or anti-proliferative effect is context dependent and the underlying mechanisms still unclear (Cui *et al*, 2016; Hu *et al*, 2021; Lazzari & Meroni, 2016; Wang *et al*, 2018; Zhao *et al*, 2018). Notably, it was reported that *Trim32* deficiency in primary myoblasts led to G0/G1 cell cycle arrest and premature senescence under growth conditions, a phenomenon that we do not observe in our C2C12 model (Kudryashova *et al*., 2012; Servian-Morilla *et al*., 2019). It is likely that this inconsistency is due to the fact that those reports referred to senescence of primary satellite/Muscle stem cells (MuSCs), while our cellular system represents a myoblasts population at a later commitment stage. This fact suggests that TRIM32 might be involved at different phases of muscular differentiation, with different impacts resulting from its deficiency ultimately leading to defective myogenesis. Indeed, we observed MyoD reduction as well as variation of global transcriptome also before the differentiation stimulus, which will need further studies.

Further investigation into the underlying mechanism revealed a Trim32-c-Myc axis in regulating cell cycle exit. High levels of c-Myc promote myoblast proliferation, however, cell cycle exit is a pre-requisite for the onset of differentiation (Buttitta & Edgar, 2007). This is achieved by rapid down-regulation of pro-proliferative factors among which is c-Myc (Bretones *et al*, 2015; Kolly *et al*, 2005). Previous studies have shown Trim32–c-Myc interaction in the context of neurogenesis, and that Trim32-mediated c-Myc ubiquitination and degradation are necessary for neurogenic differentiation (Nicklas *et al*., 2019; Schwamborn *et al*., 2009). Likewise, in the context of myogenesis c-Myc was also shown to be significantly ubiquitinated and downregulated upon overexpression of Trim32, while a Trim32 mutant (Cys24Ala) lacking E3 ligase activity failed to affect c-Myc levels and impaired the differentiation process (Nicklas *et al*., 2012; Schwamborn *et al*., 2009). However, straightforward evidence that c-Myc is a substrate of Trim32 remains lacking. Consistent with this earlier view, we found indeed sustained c-Myc expression in *Trim32* KO clones at the onset of differentiation. Surprisingly though, c-Myc protein degradation did not show significant difference in *Trim32* KO and WT clones. Instead, c-Myc mRNA stability was increased in KO cells suggesting that Trim32 regulates c-Myc expression by regulating turnover of its transcript rather than protein ubiquitination. The inconsistency with previous data on Trim32-mediated c-Myc ubiquitination and degradation may be due to the specific cell context and/or to the narrow time window we investigated, just after the switch from proliferating to differentiating conditions, where c-Myc abrupt reduction is needed to prompt the myogenesis programme. Regarding the effect on c-Myc mRNA, Trim32’s involvement in RNA regulation has been reported, particularly on microRNAs (Schwamborn *et al*., 2009), but whether Trim32 directly interacts with c-Myc mRNA or acts through intermediaries such as microRNAs or other proteins, and whether Trim32 E3 ligase activity is required, remains to be determined.

Functionally, we demonstrated that c-Myc contributes to the impaired myogenesis observed in *Trim32* KO clones, however, it is clearly not the only factor regulated within the Trim32-mediated myogenic network. While downregulating c-Myc at the onset of differentiation restores Myogenin expression in nascent myotubes, this intervention is insufficient, at least within our experimental timeframe, to fully normalize the differentiation index or full myotube morphology. Two possible non-exclusive explanations may account for this observation. On one side, c-Myc may influence early stages of myogenesis, such as myoblast proliferation and initial myotube formation, but it may not contribute significantly to later events such as myotube hypertrophy or fusion between existing myotubes and myocytes. This hypothesis is supported by recent work showing that c-Myc is dispensable for muscle fiber hypertrophy but essential for normal muscle stem cell (MuSC) function (Ham *et al*, 2025). On the other hand, the impaired myogenesis of Trim32 KO clones may not result solely from accumulated c-Myc levels. Indeed, Trim32 is known to regulate multiple molecular pathways, and its loss may affect a network of downstream targets beyond c-Myc as also mentioned above. Only in the context of regulation of cell cycle progression, which must be inhibited in order to allow the differentiation program to occur, Trim32 was for instance shown to promote ubiquitination and degradation of NDRG2, and in absence of Trim32 accumulation of NDRG2 resulted in sustained proliferation and delayed cell cycle withdrawal (Mokhonova *et al*., 2015). Thus, Trim32 may be involved at various levels of myogenic regulatory pathways, and these different Trim32-dependent mechanisms likely converge to produce the myogenic deficits characteristic of LGMDR8 patients.

In summary, our study demonstrates that the absence of Trim32 disrupts early phases of myogenesis in a cellular system. Further, we identified a novel mechanism through which Trim32 regulates the transition from proliferation to differentiation by modulating c-Myc mRNA stability, revealing an unrecognized pathogenic mechanism that might highlight potential novel therapeutic approaches for future treatment of LGMDR8.

## Methods

### Cell culture

C2C12, an immortalized murine myoblast cell line, was a kind gift of Prof. D’Andrea, University of Trieste. Cells were maintained in culture in growth medium (GM): DMEM (Euroclone, ECM0101L), 20% foetal bovine serum (FBS, Gibco, 10270), 4 mM L-glutamine (Euroclone, ECB3000D), 100 units/mL penicillin/100 μg/mL streptomycin (Gibco, 15104). Cells were maintained at 37°C in 5% CO_2_ in a humidified incubator.

For myogenesis induction, cells are seeded at 50-60% density and cultured in growth medium in either 6 or 10 cm petri dishes. At confluence, GM is replaced with differentiation medium (DM): DMEM (Euroclone, ECM0101L), 2% donor horse serum (biowest, S0900), 4 mM L-glutamine (Euroclone, ECB3000D), 100 units/mL penicillin/100 μg/mL streptomycin (Gibco, 15104), counting as Day 0 (D0) of the differentiation process. The DM was changed every day before fixation or cell harvesting according to the different experiments. The differentiation index was calculated as the ratio of nuclei within MyHC-positive myotubes to the total number of nuclei.

### Generation of Trim32 Knock-out clones

*Trim32* knock-out C2C12 clones were generated by using CRISPR/Cas9 based genome editing system (OriGene KN318203D). An all-in-one vector (pCas-Guide) with a *Trim32* target gRNA and Cas9 sequence, and a donor vector (pUC-Donor-vector) carrying GFP-LoxP-Puro-LoxP (Appendix Table) were transfected into C2C12 parental cell line using either lipofectamine 3000 (Invitrogen, L3000015) or Mirus TransIT-X2 Dynamic Delivery System (Mirus, MIR6004), as per the manufacturer’s recommendations. After 3 weeks of culturing, puromycin (1µg/mL) was applied for selection. The medium was changed every 2-3 days for 10 days and individual cell clones were isolated and expanded. Genomic DNA was extracted from each clone using the PCRBIO Rapid Extract Lysis kit following manufacturer’s instructions (PCRBIOSYSTEMS, PB15.11-24). Genomic DNA was amplified with primers flanking the targeted *Trim32* region (Appendix Table) and amplicons were Sanger sequenced. Wild-type clones were selected based on *Trim32* non-mutated sequence.

### RNA extraction and RNAseq analysis

Total RNA was extracted with the TRIzol reagent (Thermo Fisher, 15596-026) following the manufacturer’s instruction and was quantified using NanoDrop 2000; for the RNAseq experiments, RNA was also qualitatively evaluated using Agilent Bioanalyzer 2100. RNA was then stored at – 80°C until use.

For the RNAseq experiments, for each of the WT and KO samples at D0 and D3 of the differentiation process, total RNA with an RNA Integrity Number (RIN) higher than 9 was employed. For each sample, libraries were generated starting from 500 ng of total RNA by Illumina Stranded mRNA Prep, according to the manufacturer’s protocol. The libraries were quantified using Qubit dsDNA BR Assay Kit on Qubit 2.0 Fluorometer and checked for dimension with DNA 1000 Chip on Bioanalyzer 2100 before pooling. The final sequencing pool was quantified by qPCR before sequencing. Sequencing was performed on Novaseq 6000 sequencer according to the manufacturer’s protocol and generating, for each sample, almost 50 millions clusters of 2×150 bp paired-end reads. Raw files were subsequently quality checked with FASTQC software and sequences with low quality score or including adaptor dimers, were discarded from the analysis. The resulting set of selected reads were aligned onto the Mouse genome using STAR version 2.7.3 (Dobin & Gingeras, 2015) using GRCm39 Genome Assembly and Gencode.v28 as gene definition. The resulting Mapped reads were used as input for feature Counts functions of Rsubread packages and used as Genes counts for Differential expression analysis using Deseq2 package (Love *et al*, 2014). Differentially expressed genes (DEGs) were selected for |log2(FC)| −1 or 1 and corrected P value 0.05 and used as input to perform pathway enrichment analysis by IPA system (Ingenuity® Systems). Gene set enrichment analysis was conducted using the rotation gene set test function within the edgeR package (v 3.38).

The raw sequencing data generated in this study have been deposited in the NCBI Sequence Read Archive (SRA) under the accession number PRJNA1280561.

### Real Time-qPCR

cDNA was reverse transcribed from 1μg of RNA using SuperScript™ II Reverse Transcriptase (Invitrogen, 18064014), according to the manufacturer’s instructions, and cDNA stored at −20°C. RT-PCR was performed using 1/40^th^ of the generated cDNA, SensiFAST^TM^ Probe Kits (Meridian Bioscience, BIO-98020) and 200 nM of forward and reverse primers, in a final volume of 20 μL. c-Myc primers were designed to span exons. c-Myc targeted genes and Gapdh primers were reported in (Cairo *et al*., 2005). RT-PCR was performed on an CFX CONNECT SYSTEM real-time PCR machine (Bio-Rad), in technical triplicates. Gene expression changes were calculated using the 2^-ΔΔCt^ method. See Appendix Table for primer sequences.

### Treatments and transfections

Time course experiments to analyze c-Myc protein and c-Myc mRNA stability were performed by treating the cells with 25 µg/mL cycloheximide (Thermo Fisher, J66901) and 5µg/mL Actinomycin D (Sigma-Aldrich, A1410), respectively. The time of treatment is indicated in the results section.

Plasmid DNA harboring Flag-c-Myc (Appendix Table), already available in the lab, was transfected in C2C12 cells with Lipofectamine 3000 (Invitrogen, L3000015) following manufacturer’s instructions. Cells were harvested 24 hours (D0) and 8 days (D8) after induction of differentiation for Western blot analyses.

For silencing experiments, siRNA (Dharmacon, FE5L040813000010 and FE5L0018101020) and DharmaFECT reagent (Dharmacon, FE5T200102) were each diluted, combined and incubated according to the manufacturer’s instructions. The transfection mixture was brought to the appropriate volume and added at a final concentration of the siRNA of 25 nM. Cells were harvested 24 hours post-transfection in growth medium (D0), as well as at various time points after differentiation induction for RNA and protein analysis.

### Immunofluorescence Experiments

C2C12 cells were seeded on glass coverslips at a density of 7.5 x 10^4^ cells/mL. Coverslips were collected at different time points, washed twice in PBS (137mM NaCl, 2.7 mM KCl, 10 mM Na_2_HPO_4_, 1.8 mM KH_2_PO_4_, pH 7.4) and fixed in 4% paraformaldehyde (PFA) in PBS for 15 minutes at room temperature, or fixed in methanol for 10 minutes on ice (for myosin heavy chain stain). Cells were then rinsed in PBS and blocked with 5% BSA (VWR Life science, 0332) in PBS-Triton X-100 (0.3%) for 1 hour at room temperature. Coverslips were incubated with primary antibodies, diluted in 1% BSA in PBS-Triton X-100 (0.3%), overnight at 4°C. Staining was obtained after incubation for 1 hour at room temperature with a Cy3-conjugated anti-rabbit antibody (1:200 dilution, #AP132C, Millipore) or FITC-conjugated anti mouse antibody (1:200 dilution, #F0479, DAKO). Coverslips were mounted with Vectashield mounting medium plus DAPI (Vector Laboratories, H-1200) and cells were imaged with epifluorescence microscopy: 20x magnification images were used to calculate differentiation index and percentage of pHH3 positive cells, while 40x and 63x magnification images were used for more detailed analyses, through ImageJ software (NIH, Bethesda, USA). See Appendix Table for antibodies information.

### Western Blot

Cells were lysed either in RIPA buffer (50 mM Tris-HCl pH 8, 0.3% SDS, 150 mM NaCl, 0.5% Sodium Deoxycholate, 1% NP-40, 1mM PMSF and 1x Protease Inhibitor Cocktail (Sigma-Aldrich, P8340) or Myosin extraction buffer (300 mM NaCl, 0.1M NaH_2_PO_4_, 0.05M Na_2_HPO_4_, 0.01M Na_4_P_2_O_7_, 1mM MgCl_2_, 10mM EDTA, 1mM DTT, 1mM PMSF and 1x Protease Inhibitor Cocktail). Samples were sonicated 5 seconds x 3 pulses. The insoluble fraction was removed by centrifugation at 20,000 x *g* for 10 minutes at 4°C. Protein quantification was carried on with Bradford assay (Bio-Rad, #500-0006) following the manufacturer’s instructions. Proteins were boiled at 95°C for 5 minutes in 1x Laemmli buffer supplemented with 5% β-mercaptoethanol, and loaded on 7.5%, 10%, or 12% polyacrylamide gels (PAGE), depending on the resolution required, and blotted on PVDF membranes (Millipore, Immobilon-P, IPVH-00010) overnight at 4°C. Membranes were then blocked in 5% Nonfat dry milk (Cell Signaling Technology, 9999S) in TBS 1x (150 mM NaCl, 10 mM Tris pH 7.4) with 0.1% Tween (TBST), for 1 hour at room temperature. Primary antibodies were diluted in 5% Nonfat dry milk in TBST or 5% BSA in TBST and incubated for 2 hours at room temperature or overnight at 4°C. Secondary antibodies were incubated for 1 hour at room temperature, diluted in 5% Nonfat dry milk or BSA in TBST. See Appendix Table for antibodies information. For detection, membranes were incubated with HRP substrate (ECL Pierce, Invitrogen, 32106) for 1 minute, and images acquired with ChemiDoc MP Touch Imaging System (Bio-Rad). Relative bands intensities were analyzed by Image Lab software (Bio-Rad).

### MTT assay

MTT (Sigma-Aldrich, M2128) was used to determine cells metabolic activity. Cells were seeded in 96-well plates at a concentration of 6000 cells/well in 200 µL growth medium, in technical triplicate. 24 hours after seeding (D0), 10 µL MTT solution (5mg/mL in PBS; 0.5%) was added to each well and cells were kept incubated at 37°C for 4 hours. After incubation, the medium was removed and 150 µL DMSO (Sigma-Aldrich, 154938) was added to each well to dissolve the formazan crystals. The absorbance was read by NanoQuant (Infinite M200PRO) at OD=590 nm. The experiment was carried out with cells at Day 1 (D1), D2 and D3 of differentiation.

### Cell cycle and proliferation analysis by flow cytometry

For cell cycle analysis, 7.5 x 10^5^ cells were seeded in 10 cm dishes and allowed to adhere overnight in growth medium. The next day, cells were washed twice with PBS, trypsinized and collected in a 15 mL centrifuge tube (D0). Samples were spun down at 125 x *g* for 5 minutes and the supernatant was discarded and the cells were resuspended in 500 µL PBS. 1 x 10^6^ cells were counted and dropped into 1 mL of EtOH 70%, vortexed, and stored at 4°C. The same process was carried out with cells grown for 48 hours in growth medium or after induction of differentiation.

Before staining, cells were diluted by adding 1 mL PBS, and centrifuged at 850 x *g* at 4°C for 5 minutes. The residual EtOH was removed by rinsing with PBS twice, and cells were then resuspended in 1 mL PBS and stored at 4°C for 1 hour. Afterwards, the PBS was discarded by centrifugation, and cells were resuspended in propidium iodide (PI) staining solution (10μg/mL final concentration) (Sigma-Aldrich, P-4170), incubated overnight at 4 °C, and analyzed using a flow cytometer.

For Carboxyfluorescein Diacetate Succinimidyl Ester (CFSE) tracing cell proliferation, cells were counted and divided into two groups: cells without CFSE for background noise at time 0 and cells stained with CFSE for proliferation analysis; an aliquot of CFSE-stained cells was used for fluorescence intensity analysis at time 0 (starting generation), the remaining fluorescent cells were seeded in growth medium for 48 hours before being collected and analysed for green fluorescence. Staining with the viable probe was performed as follows: CFSE 5 µM was used to label 1×10^6^ cells. During the first 10 minutes, CFSE-stained cells were kept in PBS buffer; then the volume was doubled with growth medium to reach the final volume of 1 mL and incubated for another 5 minutes (all incubations at room temperature). Cells were centrifuged and rinsed with PBS to remove excess CFSE. After the washing step, cells were resuspended for flow cytometric analysis or for culturing. All steps in the presence of CFSE were performed in the dark. PI solution (10 µg/mL final concentration) was added to all samples immediately prior the flow cytometry acquisition and PI-positive cells were excluded from analysis to assess proliferation for undamaged cells only.

All Flow cytometry assays were performed using an Attune NxT instrument (Thermo Fisher Scientific) endowed with acoustic focusing technology, equipped with blue laser (488 nm, 50 mW) and standard optical bench configuration (4 color channels). After recording at least 20,000 events per run at flow rates of 100 μl/min to avoid clumping, data were saved as FCS files and subjected to software analysis using FCS express V7 De Novo Software V7 with the functions “Multicycle” for cell cycle analysis and “Proliferation fit” for cell division analysis.

### Co-Immunoprecipitation

Cells were lysed on ice for 30 minutes in ice-cold modified RIPA buffer (50 mM Tris-HCl pH 8, 150 mM NaCl, 1mM EDTA, 1% NP-40, 0.25% Sodium Deoxycholate) supplemented with 1 mM PMSF, 1 mM Na_3_VO_4_, 1 mM NaF, and 1x Protease Inhibitor Cocktail (Sigma-Aldrich, P8340). Lysates were centrifuged at 20,000 x *g* for 15 min at 4°C. Protein concentration was determined by Bradford assay (Bio-Rad, 500-0006). Five percent of each lysate was kept as input sample. Protein A agarose beads (Roche, 11719408001) were equilibrated by washing twice in modified RIPA buffer followed by centrifugation at 850 x *g* for 3 minutes at 4°C. Equal amounts of equilibrated beads were added to each tube, and samples were incubated on a rotor wheel at 4°C for 1 hour to remove the nonspecifically binding proteins. After centrifugation at 850 x *g* for 5 minutes at 4°C, 1 µg of antibody (Appendix Table) was added to the supernatant lysates and samples were incubated on a rotor wheel at 4 °C overnight. The following day, equal amounts of equilibrated beads were added into lysates, and samples were incubated on a rotor wheel at 4 °C for 3 hours to pull down the targeted proteins. The beads were then washed four times with modified RIPA buffer followed by centrifugation for 3 minutes at 850 x *g* at 4°C. The supernatant was set aside as unbound sample. Immunoprecipitates were eluted by boiling at 95°C for 5 min in 2× Laemmli buffer supplemented with 5% β-mercaptoethanol.

### Nuclear/Cytosol Fractionation

In total, 5 x 10^5^ cells were seeded in 6 cm dishes and allowed to reach confluence overnight in growth medium. The next day, the growth medium was replaced with differentiation medium, and cells were cultured for additional 6 hours. Cells were harvested with trypsin-EDTA and then centrifuged at 2,300 x *g* for 5 minutes at 4°C, followed by washing with PBS. The cell pellets were processed using NE-PER Nuclear and Cytoplasmic Extraction Reagents (78833, Thermo Fisher Scientific) according to the manufacturer’s instructions.

### Statistics

All the experiments have been performed on 3 *Trim32* KO and 3 WT independent clones (biological replicates), unless otherwise indicated. Statistical analyses were conducted using a two-tailed unpaired Student’s t-test in GraphPad Prism 10. Each value represents at least three independent experiments (technical replicates), except for CFSE tracing cell proliferation, which is the result of two replicates. A *p*-value of <0.05 was considered statistically significant. P values are denoted as follows: *p□<□0.05, **p□<□0.01, ***p□<□0.001, ****p□<□0.0001.

## Appendix Table

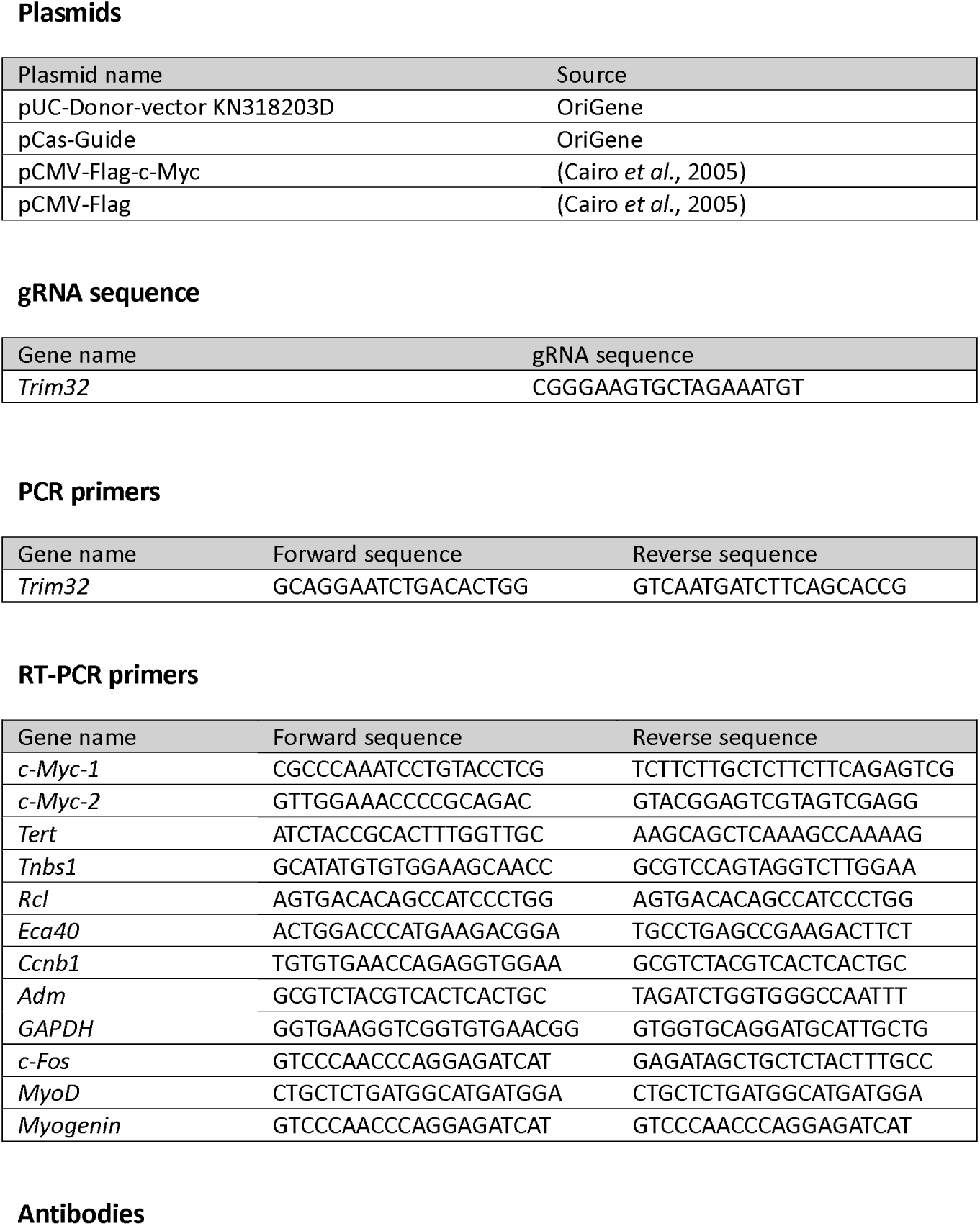
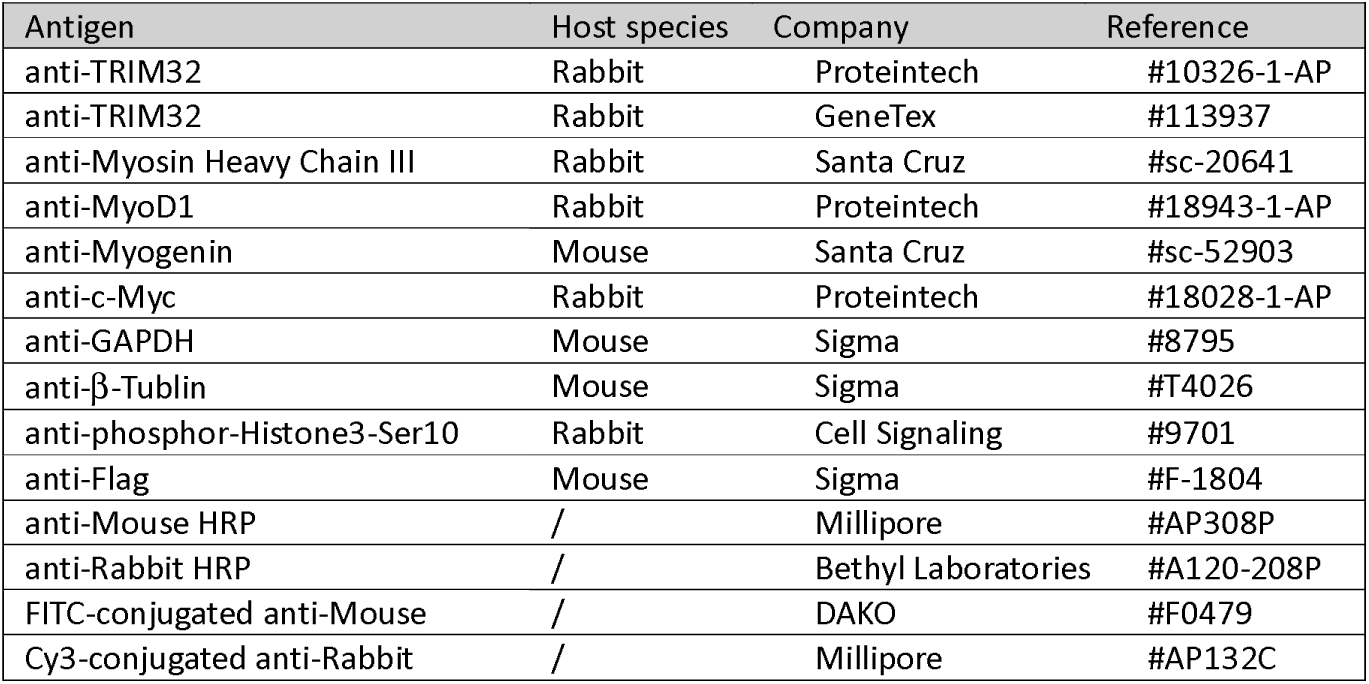

## Acknowledgements

This work has received funding from: European Union’s Horizon 2020 research and innovation programme under the Marie Skłodowska-Curie grant agreement No 813599 - TRIM-NET; AFM-Telethon, France, grant #23115; and CERIC-ERIC #20242056. L.X. was supported by Fondazione Umberto Veronesi, Italy. Meroni’s group participates to the COST Action CA20113 – ProteoCure.

Schemes used in the figures were adapted from Servier Medical Art (https://smart.servier.com/) licensed under a Creative Commons Attribution 3.0 Generic License.

## Author contributions

Lu Xiong: Investigation, Visualization, Writing – original draft, Writing – review & editing; Elisa Lazzari: Conceptualization, Investigation, Writing – original draft, Writing – review & editing; Sabrina Pacor: Investigation, Writing – review & editing; Simeone Dal Monego: Investigation; Erica Piovesan: Investigation, Writing – review & editing; Danilo Licastro: Conceptualization, Formal analysis, Writing – review & editing; Germana Meroni: Conceptualization, Supervision, Funding acquisition, Writing – original draft, Writing – review & editing.

