## Supplemental data file for "TRIM32 controls timely cell cycle exit in muscular differentiation through c-Myc down-regulation"

### Figure EV1

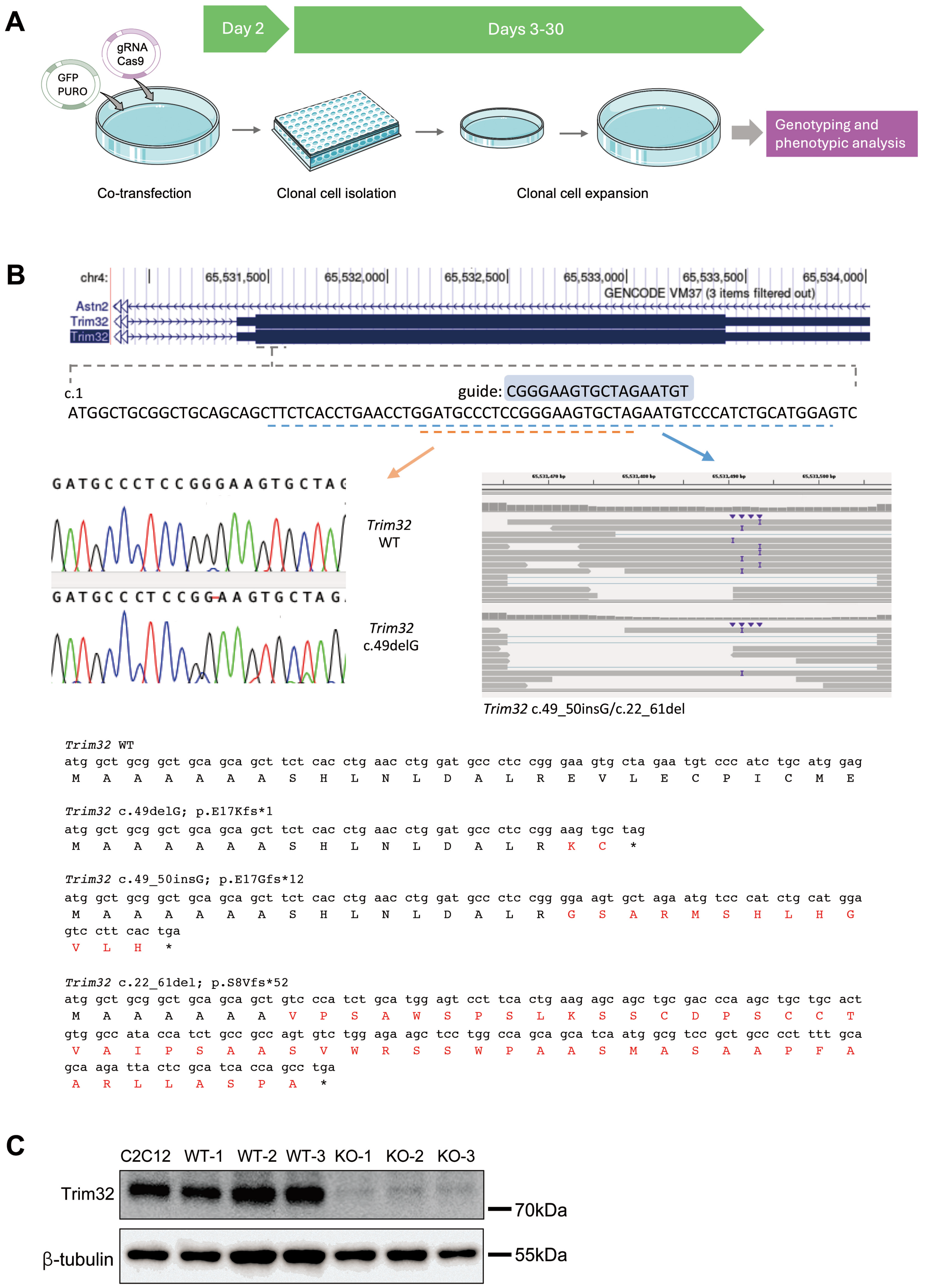


**Figure EV1 caption in the next page.**

**Figure EV1. Generation and characterization of *Trim32* KO and WT C2C12 clones. (A)** Scheme of CRISPR/Cas9 approach to generate C2C12 myoblasts *Trim32* KO clones. **(B)** Diagram illustrating the structure of the mouse *Trim32* gene and the positions of the designed gRNA. Below, on the left side, Sanger sequencing results depict the WT allele (upper electropherogram) and the c.49delG mutated allele (lower electropherogram). On the right side, RNAseq reads in the indicated genomic region are shown, confirming the compound heterozygous alleles c.22_61del, and c.49_50insG. Bottom, DNA sequence of the different mutations compared to the WT sequence showing the consequences on the encoded protein. **(C)** Representative Western blot analysis of Trim32 protein in parental C2C12 cells, three KO clones, and three WT clones (n = 3). Trim32 was detected using an antibody against the central region.

**Figure EV2**

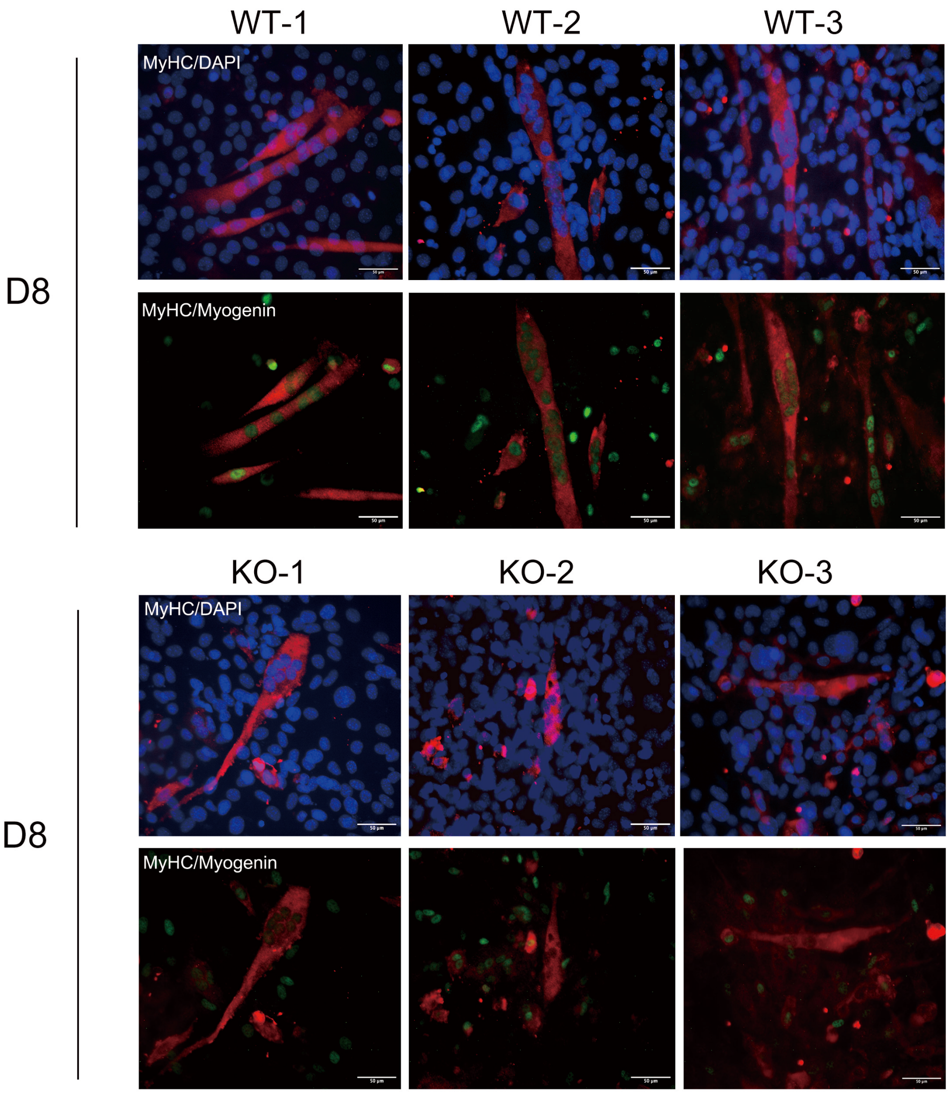


**Figure EV2.** ***Trim32*** **KO cells show impaired Myogenin expression**. Immunostaining for MyHC (red) and Myogenin (green) in WT and *Trim32* KO clones, and parental C2C12 cells at 8 days of differentiation. Nuclei were counterstained with DAPI (blue) (Scale bar = 50 µm; magnification 40x; n = 3).

**Figure EV3**

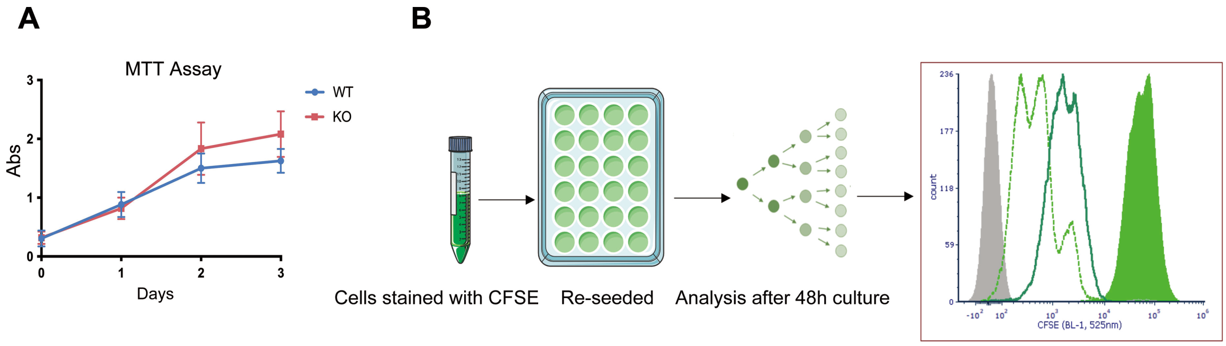


**Figure EV3. *Trim32* KO cells show higher proliferation rate compared to WT clones. (A)** Graph representing MTT assay performed to evaluate cell metabolic activity. Cells were cultured in DM for the indicated time points and absorbance at 570 nm was measured after MTT addition. **(B)** Measurement of WT and *Trim32* KO clones proliferation by CFSE (Carboxyfluorescein Diacetate Succinimidyl Ester) dilution. Cells were labeled with CFSE and cultured in DM for 48 hours. The exponential axis shows decreased fluorescence after 2 days of culturing, displaying the different proliferating rate in WT and *Trim32* KO clones. Grey area: background noise; Green area: Start generation; Dark green line: WT cells; Light green dotted line: KO cells.

### Figure EV4

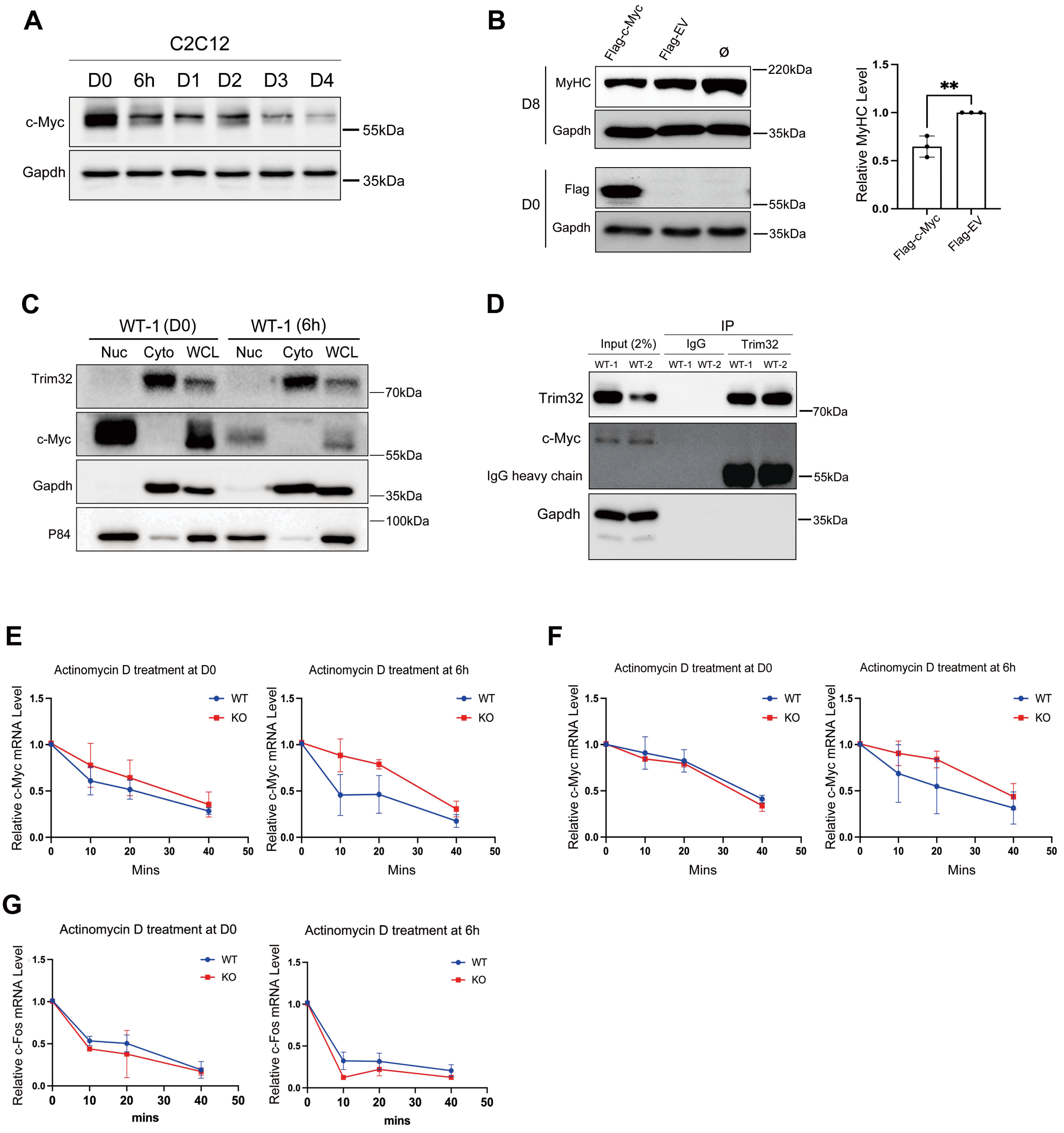


**Figure EV4. Trim32-mediated c-myc downregulation occurs at mRNA and not protein level. (A)** Western blot analysis of c-Myc protein levels in parental C2C12 at D0, and at different time points during differentiation (6 h, D1, 2, 3, and 4). Gapdh was used as loading control. **(B)** Western blot analysis of MyHC protein levels upon exogenous overexpression of Flag-c-Myc and Flag-Empty Vector (Flag-EV) in C2C12 parental cells (ø: Non-transfected). The expression of Flag-c-Myc was detected after 24 hours (D0) post-transfection and MyHC was detected at 8 days of differentiation. Gapdh was used as loading control. The accompanying graph shows the quantification from three independent replicates (mean ± SD; n=3; unpaired t-test, $^{**}P<0.01$). **(C)** Nuclear (Nuc) and cytosolic (Cyto) fractions from WT clones were analyzed by western blot for Trim32 and c-Myc. Gapdh and p84 served as markers for cytoplasmic and nuclear fractions, respectively. WCL: whole cell lysate. **(D)** Immunoprecipitation (IP) with a Trim32 antibody followed by Western blot analysis to detect Trim32 and c-Myc in representative WT clones after 6 hours of culture in DM. IP with IgG was used as control. **(E-F)** Graphs showing the stability of c-Myc mRNA determined by Actinomycin D in WT and *Trim32* KO clones. Cells were cultured in either GM or DM for 6h, followed by treatment with Actinomycin D for the indicated time points. Relative RNA levels were measured by real-time RT-PCR and normalized to Gapdh (mean ± SD; n = 3). **E**) Same as for the experiment in Figure 5E but using twice the amount of cDNA. **F)** Same as in E) but using c-Myc primers-2. **G**) The stability of c-Fos mRNA was measured as control.
